# Linking automated image analysis to ecological inference: high-throughput monitoring of soil fauna

**DOI:** 10.64898/2026.06.16.732537

**Authors:** Hadrien Hendrikx, Emma Belaud, Francois Postic, Morgan Scalabrino, Marion Lebeau, Guerric Le Maire, Christophe Jourdan, Philippe Gallet, Mickael Hedde

## Abstract

1 - Automated in situ sensors – e.g., buried scanners – are transforming biodiversity monitoring by generating data at spatio-temporal resolutions unattainable through traditional sampling, including in cryptic environments such as soil that have remained largely inaccessible to existing methods. However, extracting ecologically meaningful information from these data streams requires substantial image processing effort that currently constitutes a critical bottleneck, particularly when the signal-to-noise ratio is low and annotated training data are scarce.
2 - Standard end-to-end deep learning detection pipelines offer unsatisfactory results due to the lack of training data and heterogeneity of the taxa of interest. We explore the potential of combining traditional computer vision algorithms with state-of-the-art deep learning models to build an efficient raw data processing pipelines from limited annotation effort. Specifically, based on the observation that the background barely changes, we focus on the differences between two consecutive images to turn the initial detection problem (with very low signal) into a simpler classification problem, which we solve by fine-tuning foundation models on limited annotated data.
3 - Our approach significantly reduces the annotation effort, allowing us to release an open dataset with about 600 soil scans and more than 8 000 labeled invertebrate occurrences across nine taxa. Using this dataset to train our models, we obtained population count estimates with relative errors ranging from 10% to 61% across taxa over a three-month period. Ecological validation through a land-use stability analysis showed full directional congruence between automated and expert-annotated classifications across all nine taxa examined, with effect-size discrepancies proportional to per-taxon classification accuracy.
4 - These results demonstrate that combining domain-specific heuristics with fine-tuned foundation models provides an effective and data-efficient strategy for automating ecological image processing workflows in low-signal, data-scarce contexts. The validated pipeline removes the manual annotation bottleneck that has historically limited scanner-based soil monitoring to short observational windows and restricted taxonomic scope, opening the way for continuous, large-scale tracking of soil invertebrate community dynamics at resolutions previously unachievable.

## 1 Introduction

### 1.1 The rise of automated monitoring and machine learning in ecology

Ecology is undergoing a rapid technological transformation, with the growing deployment of automated sensors generating high-resolution data across space and time [Allan et al., 2018, Lahoz-Monfort and Magrath, 2021, Besson et al., 2022, Kerry et al., 2022, Stephenson, 2020, Turner, 2014]. These systems, including camera traps and other imaging devices, have become key tools for biodiversity monitoring, enabling continuous observation of ecological communities at unprecedented temporal resolution [Fisher, 2023, Oliver et al., 2023].

However, these advances come with a fundamental trade-off: increased data volume and complexity require automated processing pipelines to extract relevant ecological information [Besson et al., 2022, Kitzes et al., 2026, Tuia et al., 2026]. Over the last decade, machine learning – and computer vision in particular – has achieved major progress in tasks such as object detection, classification, and segmentation [Stevens et al., 2024, Nguyen et al., 2024]. The emergence of pretrained foundation models has further reduced the need for large annotated datasets, enabling transfer learning across tasks and domains [Radford et al., 2021, Oquab et al., 2023, Woo et al., 2023, Siméoni et al., 2025].

These approaches are increasingly integrated into automated biodiversity monitoring systems [Chimienti et al., 2026, Besson et al., 2022], but their application remains challenging in ecological contexts characterized by low signal-to-noise ratios, strong domain shifts, and limited training data. Beyond these technical difficulties, downstream use in ecological research incurs additional challenges, as globally efficient but biased models may lead to wrong ecological conclusions.

### 1.2 Soil ecosystems: a frontier for automated ecological observation

Soil ecosystems represent a particularly challenging and underexplored domain for automated monitoring. Although it hosts nearly a quarter of the Earth’s biodiversity for all or part of their life cycle, current methods for studying these organisms are highly restrictive [Anthony et al., 2023, Decaëns et al., 2006, Geisen et al., 2019]. As soil is an opaque matrix that creates a significant physical barrier to study, in situ observations and sampling of edaphic organisms are challenging [Erktan et al., 2020]. Thus, existing approaches – such as soil cores, hand sorting, or pitfall trapping – provide temporally sparse and spatially aggregated snapshots of communities.

As a consequence, key ecological processes, including short-term population dynamics and responses to environmental variability, remain poorly resolved in soil systems. This limitation restricts our ability to understand how soil biodiversity responds to land use and environmental change, and to integrate soil organisms into broader ecological monitoring frameworks.

Recent developments in in situ imaging systems, such as buried optical scanners [Belaud et al., 2024], provide new opportunities to address these limitations by generating continuous image time series of soil fauna. These systems enable non-destructive observation of organisms within the soil matrix at high temporal resolution, potentially transforming how soil biodiversity is monitored. While the promises are great, manually processing all images is infeasible over long periods of time, so that scanners alone are not sufficient to actually perform this deep transformation of soil biodiversity monitoring.

### 1.3 Automated pipelines as tools for ecological inference

A natural answer is to build computer vision pipelines to automatically process scanner images. Yet, despite the recent advances in computer vision, transforming raw sensor data into ecologically meaningful variables remains a major challenge. This challenge is rooted in a fundamental principle of ecology: observed data are often imperfect proxies of underlying ecological states due to detection and observation errors. Whether based on traditional trapping or modern automated imaging, ecological data are shaped by the interaction between true biological processes and the observation systems used to capture them. Importantly, automated pipelines do not simply process data – they define how observations relate to ecological states. In this sense, they can be viewed as implicit observation models, linking raw data to ecological variables.

These automated pipelines typically involve multiple steps, including detection of candidate objects, classification into taxonomic groups, and aggregation into ecological metrics such as counts or activity indices [Zurowietz et al., 2018]. Each of these steps introduces specific sources of uncertainty, including false positives, false negatives, and misclassification errors. In this context, the limiting factor in biodiversity monitoring is no longer data acquisition per se, but the ability to extract ecologically meaningful information from these complex and often poorly characterized observation processes. Indeed, errors introduced at any stage may propagate through the workflow and ultimately affect ecological inference, potentially leading to biased conclusions if not explicitly evaluated [Wyatt et al., 2026]. Moreover, high technical performance (e.g., in detection or classification metrics) does not necessarily guarantee ecological reliability, particularly when model errors are unevenly distributed across taxa or environmental conditions. This creates a potential gap between advances in data processing and the requirements of robust ecological inference.

Despite this, most studies in automated ecological monitoring validate their approach on technical aspects, with limited assessment of their actual implications for ecological interpretation. Developing efficient automated biodiversity data processing pipelines that are also shown to be ecologically reliable is absolutely critical, and requires close collaboration between ecology and computer vision [Weinstein, 2018, Besson et al., 2022, Kitzes et al., 2026, Cowans et al., 2026].

### 1.4 Objectives of this study

Here, we present a collaborative effort between soil ecologists and computer scientists to develop and evaluate an automated pipeline for processing data from scanner-based imaging sensors, delivering on the promise of large-scale, in situ soil fauna monitoring enabled by scanning technologies. Our objective is not only to develop an efficient processing workflow, but also to assess its implications for ecological inference. Specifically, we:

1. release an open dataset of annotated soil invertebrate images spanning multiple taxa and acquisition conditions, providing a resource for automated biodiversity monitoring^1^;
2. develop a processing pipeline combining heuristic detection and deep learning classification to extract taxon-resolved observations from raw image sequences^2^;
3. evaluate the ecological validity of the pipeline by comparing automated and expert-derived estimates of population dynamics under contrasting land-use conditions;
4. examine how classification performance relates to ecological metrics, including population counts and temporal variability.

By explicitly linking automated data processing to ecological interpretation, this study contributes to bridging the gap between advances in computer vision and the requirements of robust biodiversity monitoring, particularly in environments where direct observation remains challenging.

## 2 Materials and methods

### 2.1 Raw data: soil images from buried scanners

In situ raw data were acquired using an innovative soil imaging system based on modified flatbed optical scanners (Epson V39 Perfection). Four units were installed at the DIAMs experimental agroforestry site (INRAE Diascope station, Mauguio, France) across two distinct land-use treatments: two units within an arboreal linear feature (mixed tree and grass strip), and two in the adjacent cultivated crop zone. Images were collected at a 6-hour temporal frequency from March 6 to May 19, 2024 in A4 format (21×29.7cm) at 1200 dpi, leading to 10200×14032 RGB images. Further details about the acquisition methods can be found in Belaud et al. [2024]. Sample scanner images are shown in the Figure 6 (in the appendix).

### 2.2 Detection of boxes that likely contain invertebrates through image differences

Counting invertebrates directly from raw images is impractical: the size ratio between individual invertebrates (from approximately 100 pixels) and full scans (10^8^ pixels) reaches 10^6^, making manual labelling extremely time-consuming and error-prone. We estimate that a trained annotator requires approximately 30 minutes per scan, precluding real-time annotation of four scanners. Existing studies have consequently focused on a limited number of taxa and individuals [Pruvost et al., 2022], or have required exceptional annotation effort without recording bounding boxes [Belaud et al., 2024], limiting their utility for training detection models. While off-the-shelf machine learning methods for detection could be used [Pruvost et al., 2022], there is very little to no data at this point to train them, leading to poor performances even on a restricted number of taxa that are rather easy to distinguish visually.

State-of-the-art detection pipelines typically consist of two components: a candidate box proposer and a box scorer [Ren et al., 2015]. Because the signal-to-noise ratio in our images is very low and labelled training data are scarce, we replace the data-driven box proposer with a heuristic based on inter-frame image differences. This substitution simplifies the learning problem (from detection to classification) and reduces annotation effort, as annotators need only inspect small candidate crops rather than full scans.

The heuristic approach exploits the structured acquisition protocol: the background is nearly stationary across 6-hours successive images, whereas most invertebrates move. Candidate regions are identified by focusing on pixels for which the radiometry change between consecutive frames. This is implemented in the pysoilscan software (available with the rest of our code), which relies on the skimage library [van der Walt et al., 2014], and proceeds in three steps (Figure 1):

1. Co-registration: images are automatically re-aligned, using (optical_flow_ilk) primitive, to correct pixel shift caused by mechanical vibration. Before this, the grey-scale images are normalized (column-wise). This is important because the scanners use ”push-broom” type of scanning diodes that may not be well intercalibrated on the white band of the scanner previous to a scan.
2. Adaptive binarisation: the pixel-wise grayscale difference between two consecutive scans, based on the red band is computed and adaptively thresholded, retaining only regions of significant change as candidate detections.
3. Region-of-interest identification: contiguous high-difference regions are merged after dilation-erosion steps and enclosed in bounding boxes using the regionprops and find_contours primitives.

**Figure 1:**
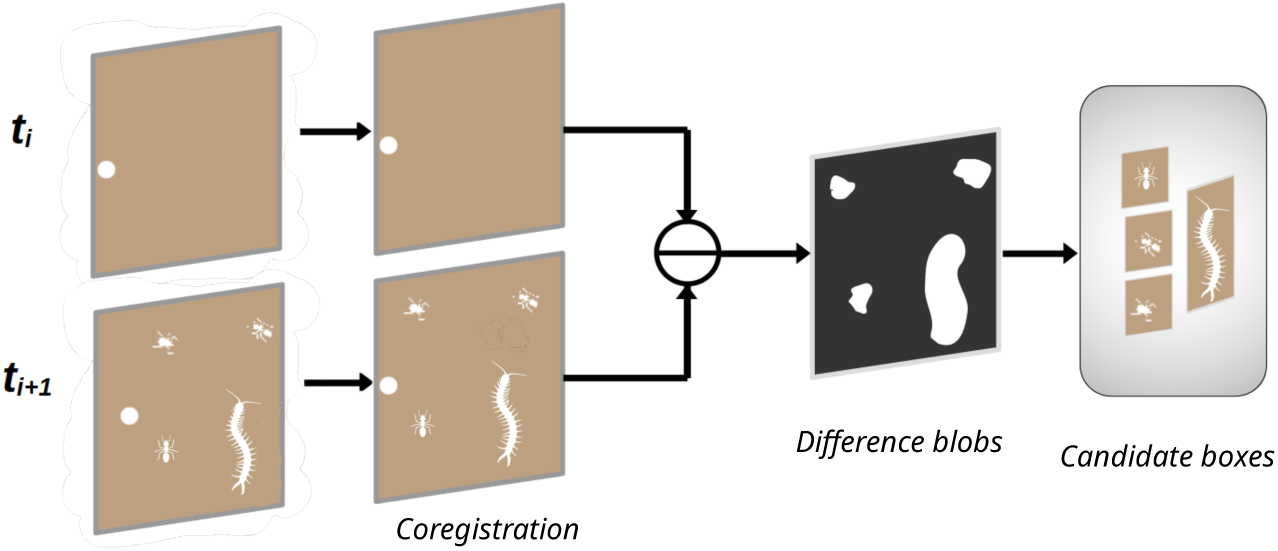
Visual representation of the steps of the heuristic detection step of our pipeline. Coregistration is a method that aligns two related images, difference blobs are computed by thresholding greyscale differences, and candidate boxes are extracted by identifying regions of interest in the difference blobs map.

We detect candidate objects using differences between successive images. This step prioritizes sensitivity and may generate false positives (e.g. noise or non-biological movement) as well as false negatives when organisms remain stationary, thus influencing the detection component of the observation process.

### 2.3 Manual annotation

Despite the high false-positive rate of the detection step (approximately 85% of extracted boxes contain no invertebrate), the reduction in search space is tremendous and makes manual annotation feasible. Soil ecologists reviewed all extracted boxes and assigned taxon labels based on the visible morphological characteristics of organisms, without predetermined taxonomic precision or class constraints. This produced a diverse set of labels spanning multiple invertebrate taxa alongside non-faunal detections (roots, fungal structures, and organic matter fragments). Rare taxa were merged into an unknown class and all non-invertebrate detections were consolidated into a background class to reduce label noise during model training. The final dataset comprises nine invertebrate classes (see Table 2). Annotation time ranged from less than ten seconds for unambiguous background boxes to several tens of seconds for invertebrate individuals requiring careful examination. This throughput remains impractical for continuous long-term monitoring at multiple sites. The resulting annotated dataset, hereafter referred to as the ’DIAMS dataset’, simultaneously enabled the ecological study reported here and served as training data for the classification models.

### 2.4 Fine-grained classification of candidate boxes into actual taxa

#### 2.4.1 Models and training

All models were initialised from pretrained foundation model checkpoints and fine-tuned on our dataset, following standard transfer-learning practice. The primary architecture was ConvNeXtV2-Base [Woo et al., 2023] (88 million parameters; checkpoint convnextv2_base.fcmae_ft_in22k_in1k from the timm library), with a fully connected linear classification head and softmax output on the logits (2 classes for binary, 13 for multiclass). We additionally evaluated DinoV3-Base [Siméoni et al., 2025], a Vision Transformer (checkpoint vit_base_patch16_dinov3.lvd1689m from the timm), as an alternative backbone. Input boxes were resized to 224 *×* 224 (ConvNeXtV2) or 256 *×* 256 (DinoV3). Training augmentations comprised random rotations, horizontal and vertical flips, and CutMix-Mixup [Zhang et al., 2017, Yun et al., 2019]. Models were fine-tuned for 30 epochs with a batch size of 64 using the AdamW optimiser [Loshchilov and Hutter, 2017], a peak learning rate of 10*^−^*^3^, five linear warm-up epochs from 10*^−^*^4^, and cosine annealing thereafter. Final performance was computed on an exponential moving average of model weights (decay = 0.999).

Model selection used scanner-stratified cross-validation to prevent data leakage: background texture is scanner-specific, so random or temporal splits would inflate validation performance. One scanner with a markedly different data distribution was retained in all training folds; three folds were obtained by leaving out each of the remaining scanners in turn. Performances were evaluated using mean Average Precision (mAP), which corresponds to the area under the precision-recall curve. Metrics reported (both mAP and counts) are validation metrics, meaning that they were used to select model hyperparameters – thus possibly slightly inflating performances, though hyperparameter tuning remained limited. We report the standard deviation computed across the different folds, i.e., when varying the scanners used for training and evaluation.

#### 2.4.2 Binary filtering and fine-grained classification

We evaluated three classification strategies: (i) binary classification to discriminate invertebrates from background, (ii) a two-stage approach combining binary pre-filtering with a dedicated multiclass classifier trained on a balanced dataset, and (iii) direct multiclass classification to assign each box to a taxon.

Binary classification. Binary models were trained on the full dataset with all invertebrate classes merged into a single positive class, and were designed to discard the large fraction of background boxes generated by the candidate box detection stage. Performance was assessed using two mAP formulations: micro-average mAP, treating all invertebrates as a single class (with invertebrate scores summed across taxa for multiclass models), and binary macro-average mAP, computed as the mean of per-taxon binary mAPs after discarding all other invertebrate examples. In this second approach, for a given class, all other invertebrate examples (excluding the background and the class of interest) were discarded; the *mAP* was then computed on the resulting binary dataset consisting only of images of a given invertebrate and background, and this per-class binary mAP is averaged across classes. This can be understood as an average of class-vs-background results (instead of the standard one versus-all). For the multiclass models, we used the score given by the class of interest instead of the sum of all invertebrates scores, so the models predict detections for a given class, and not a broad invertebrates class. We call this model Binary.

Precise invertebrate classification. Multiclass models were trained on a pre-filtered subset of the original dataset, obtained by only keeping background examples that are scored above a given threshold by the binary model. This reduces the proportion of background examples in training while focusing the model on the more challenging discrimination among taxa. Per-class mAP was computed in a one-versus-all fashion and macro-averaged across classes and cross-validation folds. For the two-stage model, mAP computation accounts for the two independent score thresholds (binary and multiclass stages). We refer to the output of this two-stage pipeline (binary then multiclass classification) as ConvBinMul.

Direct invertebrate classification. As a simpler baseline, multiclass classification was applied directly to the full, heavily imbalanced dataset without binary pre-filtering of the background class. We refer to the model obtained in this way using the ConvNextV2-base backbone as ConvDirect. Similarly, DinoDirect and ConvTiny refer to the models obtained respectively with DinoV3-base, ConvNextV2-Tiny backbones. Finally, ConvFrozen refers to a direct classification model with a frozen ConvNextV2-base backbone (only the classification head is trained).

### 2.5 From individual detections to population counts

Although mAP is the standard performance metric for classification, the ecologically relevant output is invertebrate abundance averaged over time. We therefore report, alongside mAP, population counts per scanner aggregated over either 7-day windows (short-term dynamics) or the full 3-month study period. Agreement between predicted and ground-truth counts was quantified as the mean relative error: for each time frame, we compute the absolute difference in counts, divided by the mean true count per time frame, averaged across scanners evaluated in the held-out validation folds.

To assess pipeline performance under a more pronounced distributional shift than that encountered across scanners within the DIAMs site, the full automated pipeline – heuristic detection followed by direct multiclass classification with ConvDirect– was trained on the DIAMs dataset, and then applied to raw scan data from Belaud et al. [2024]. This external dataset provides raw scanner images and manually derived invertebrate counts but no annotated detection boxes, meaning the pipeline was deployed in its intended operational mode: processing unseen raw images end-to-end without any site-specific annotation. Images were acquired from a scanner buried in a truffle oak field over three months, yielding soil conditions, background texture, and invertebrate community composition that differ substantially from the DIAMs training set. Population counts extracted by the automated pipeline were compared directly against the published manual counts, providing a stringent test of cross-site transferability under realistic deployment conditions.

Detections are aggregated into counts per time window to derive temporal activity patterns. This aggregation step may smooth short-term fluctuations and amplify persistent detection biases, influencing estimates of temporal variability.

### 2.6 Pipeline validity in an ecological use-case application

To evaluate the ecological utility and taxonomic reliability of the automated pipeline, we benchmarked it against a parallel expert-annotated analysis on the DIAMS dataset. Specifically, we tested the hypothesis that arboreal linear features in agroforestry systems (Position A) act as stable micro-refugia for soil invertebrate taxa, whereas adjacent cultivated zones (Position C) exhibit volatile, pulsed population dynamics. Population stability was defined as the inverse of temporal variability, quantified using the coefficient of variation (CV) within a 7-day rolling window applied to a 33-day continuous time series (6 March – 8 April 2024; four observations per day at 6-hour intervals). Rolling windows containing more than 5% missing observations due to sensor downtime were excluded to prevent technical artefacts from inflating apparent biological variability.

Rolling CV values are strictly positive and right-skewed; we therefore fitted Zero-Inflated Gamma (ZIG) generalised linear mixed models (GLMMs) using the glmmTMB package [McGillycuddy et al., 2025] in R. A first-order autoregressive (AR1) correlation structure was incorporated as a random effect over scanner orientation to account for temporal autocorrelation in the rolling CV series. Where the AR1 model failed to converge (non-positive-definite Hessian), a simpler random-intercept model for scanner orientation was substituted. Model goodness-of-fit was assessed using simulation-based residual diagnostics implemented in DHARMa [Hartig, 2024], testing for overdispersion, outliers, and deviations from the expected residual distribution (see Supplementary Material). Technological congruence was assessed by replicating the entire analytical workflow on data classified solely by the automated model. In order to use the full dataset and avoid data leakage, counts for one scanners are obtained by training on the other ones. Yet, the training epoch to keep was selected according to performance of the test scanners, which can lead to a slight overevaluation of the model performances. Land-use effect sizes (position C versus position A) from both pipelines were compared using a 1 : 1 identity plot, with congruence evaluated as the absolute difference in effect estimates across methods.

## 3 Results

All these results can be reproduced using our code, and our dataset.

### 3.1 Detection of boxes that likely contain invertebrates through image differences

The heuristic detection algorithm was applied to 610 scan images, yielding approximately 56k candidate boxes. The algorithm achieved high recall at the cost of low precision: approximately 85% of extracted boxes belonged to the background class. Despite this imbalance, the algorithm substantially improved the invertebrate-to-background ratio relative to raw scans. After excluding the 0.5% largest boxes (which correspond exclusively to background), the preprocessing step reduced the number of pixels requiring annotation by a factor of approximately 200. Expressed as a signal-to-noise ratio, the proportion of pixels from boxes containing invertebrates increased from 0.44% (relative to total scan pixels) to 20.9% (relative to pixels within all extracted boxes), or equivalently to approximately 15% – the proportion of non-background boxes. These estimates assume near-perfect recall. Boxes areas of invertebrate detections spanned a wide range, extending over areas from less than 10^2^ to up to 10^7^ pixels, with detection counts per image predominantly between 0 and 25 (see Appendix A.1 for more details).

More information about the dataset – including ground truth reliability and samples images – can be found in Appendix A. The full dataset is hosted on Zenodo, and can be accessed by following this link. Sample images for several invertebrates classes (as well as a root) are provided in Figure 2.

**Figure 2:**
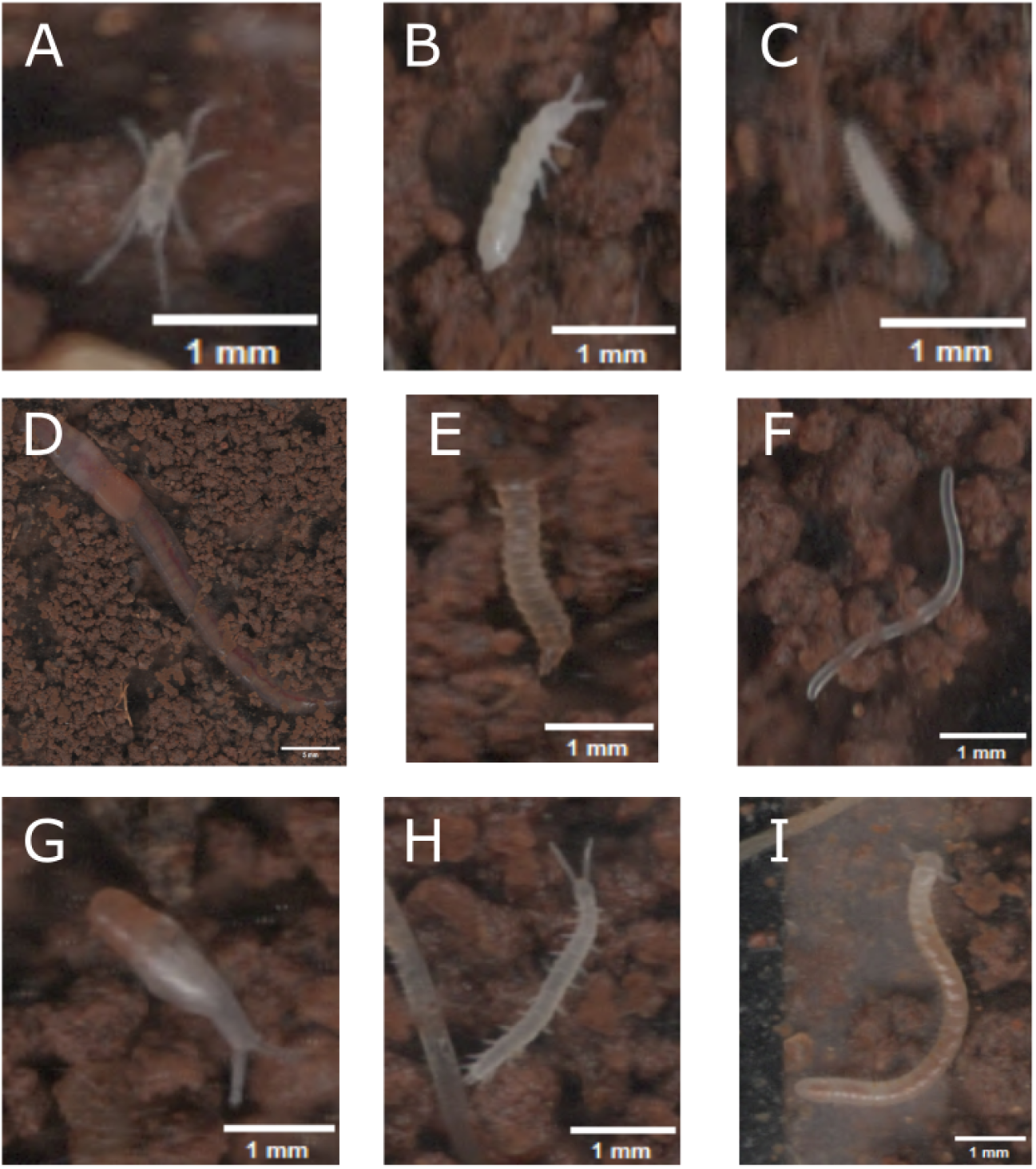
Examples of boxes detected by our image differences algorithm (one image per class of invertebrates). Base images vary in sizes, as can be seen from size of the 1mm scale. A: acari, B: collembola, C: pauropoda D: lumbricidae, E: insecta larva F: enchytraeidae, G: gastropoda, H: symphyla I: julida. More images can be found in Appendix A.

### 3.2 Fine-grained invertebrates classification and population counts

Classification performance was evaluated on two tasks: binary discrimination of invertebrates from background, and multiclass taxon assignment. Results for all model variants are reported in Table 1. Convolutional and transformer-based architectures produced near-identical results: ConvDirect and DinoDirect achieved comparable mAP across all tasks, with differences not exceeding 1.3 percentage points. The smaller

**Table 1:** Validation mean Average Precision (mAP, %) comparison in different settings (binary and multi, micro and macro-averaged) for various model choices. Binary macro-average mAP for each class is computed by removing all instances of other invertebrates classes.

| Setting | Binary | ConvFrozen | ConvDirect | ConvTiny | DinoDirect | ConvBinMul |
| --- | --- | --- | --- | --- | --- | --- |
| Binary – micro | 96.3 ± 0.4 | 91.6 ± 1.5 | 95.7 ± 0.6 | 95.7 ± 0.3 | 95.9 ± 0.6 | 96.4 ± 0.3 |
| Binary – macro | 67 ± 7.0 | 65.8 ± 2.6 | 86.3 ± 1.9 | 85.2 ± 2.6 | 85.0 ± 1.8 | 89.3 ± 1.2 |
| Multi – macro | – | 55.5 ± 1.2 | 79.1 ± 1.9 | 77.8 ± 3.0 | 77.9 ± 2.1 | 80.8 ± 1.7 |

ConvTiny yielded competitive performances (77.8% multi-macro mAP vs 79.1% for the base model) at approximately one third of the computational cost. On the other hand, training a linear head on top of a frozen backbone (ConvFrozen) resulted in a substantial performance drop across all metrics. The best performances were systematically achieved by ConvBinMul.

Aggregate metrics concealed substantial variation across taxa (Table 2), with per-class multiclass mAP ranging from 63% (Insecta larvae) to 96% (Symphyla). This variation is not limited to fine-grained classification, since binary mAP also ranged from 75% (Insecta larvae) to 97% (Collembola, Gastropoda). Count errors followed a broadly consistent pattern across taxa: relative errors were higher at the 7-day scale than over the full 3-month period, as expected given the reduced opportunity for error compensation over shorter windows. A positive association between the technical metric (classification accuracy) and the ecological signal (population counts reliability) was apparent across taxa (Figure 3, left panel), whereas the relationship between training set size and count error can be identified (in particular after removing outliers such as enchytreidae), but is weaker (Figure 3, right panel). Collembola and Gastropoda achieved both high mAP (94% and 91%, respectively) and low count errors (10–12% and 16–18% across temporal scales), while Enchytraeidae combined moderate mAP (67%) with the highest count errors of all taxa (61% and 68% at 3-month and 7-day scales). Symphyla, despite being among the rarer taxa (152 annotated individuals), achieved very high mAP (96%) and low count errors (10% and 13%).

**Table 2:**
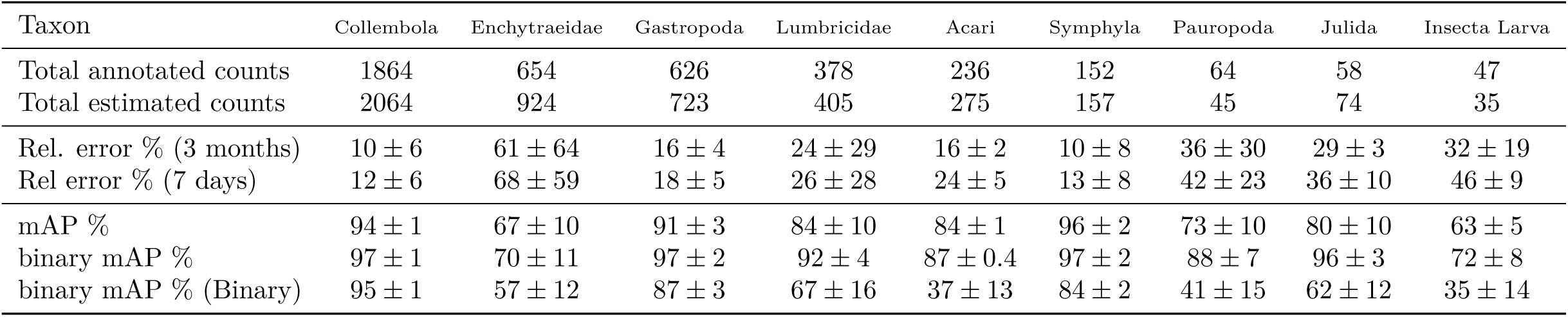
Human-annotated and model-estimated counts, relative count errors over two temporal scales, and mean Average Precision (mAP, %) per taxon for model ConvDirect. Counts are obtained by aggregating all invertebrates predictions over successive fixed time-windows of different lengths – 7 days or the whole time frame, i.e., 3 months. Relative error is obtained by computing the absolute difference between true and estimated counts for each time frame, and rescaling by the average number of true counts. Binary mAP for each class is computed by removing all instances of other invertebrates classes.

**Figure 3:**
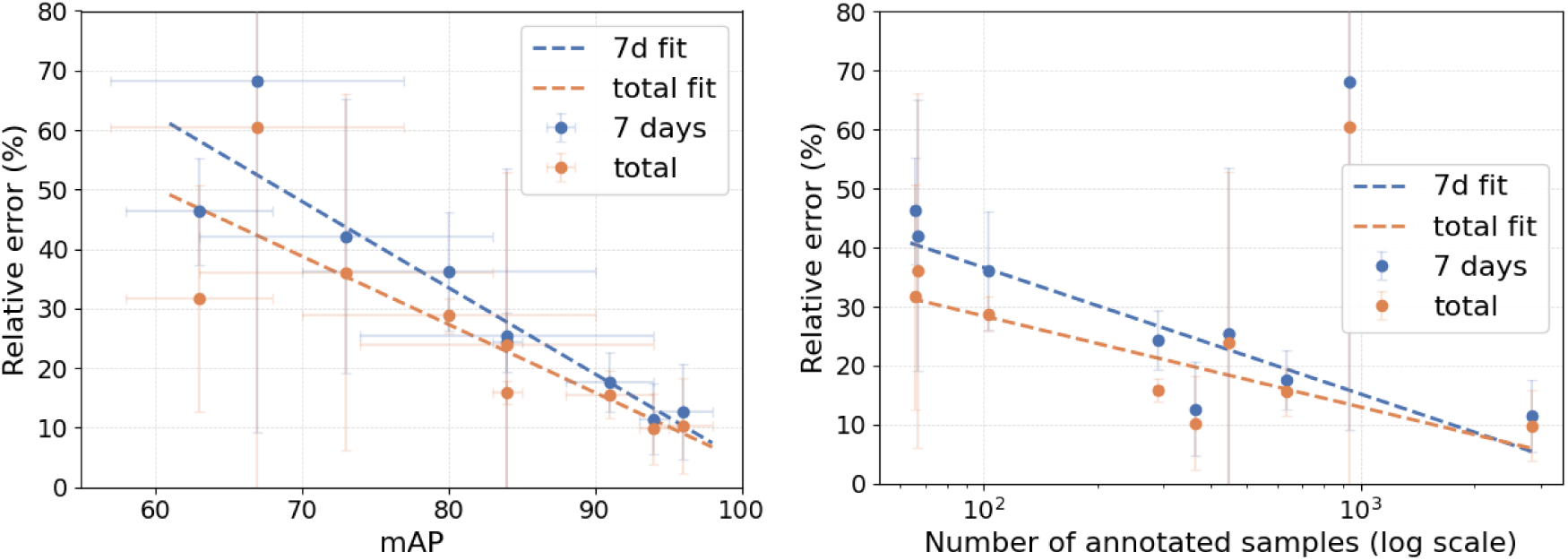
Relative count error (7-day and 3-month) as a function of mean Average Precision (left) and total number of annotated examples (right), per taxon. Counts are obtained by aggregating all invertebrates predictions over successive fixed time-windows of different lengths – 7 days or the whole time frame, i.e., 3 months, and relative error is obtained by computing the absolute difference between true and estimated counts for each time frame, and rescaling by the average number of true counts. We consider the mean number of training samples for the right figure, and exclude the uppermost outlier (enchytraiedae) from the fit.

Results from the pipeline transferability test in a contrasting context showed, on average, higher count errors than those obtained within the DIAMS site, but with marked heterogeneity across taxa (see Table 3). Collembola and Acari exhibited reasonable cross-site transferability (3-month relative errors of 7% and 18%, respectively, for ConvDirect), whereas Enchytraeidae errors remained high, consistent with their low within-site classification accuracy. Symphyla and Julida exhibited significant errors with both ConvDirect and ConvBinMul, particularly Julida, for which the 3-month relative error exceeded 390% — likely reflecting their very low abundance in the external dataset combined with a substantial shift in background composition. ConvBinMul outperformed ConvDirectfor most taxa, but this came at the cost of substantially inflated errors for some of them (e.g. Julida).

**Table 3:** Human-annotated and model-estimated counts for ConvDirect and ConvBinMul models under a pronounced testing distribution shift – relative to the training set. Models were trained on the dataset we release, and tested on data from Belaud et al. [2024], which corresponds to different scanners in a significantly different context. Counts are obtained by aggregating all invertebrates predictions over successive fixed time-windows of different lengths – 7 days or the whole time frame, i.e., 3 months.

|  | Species | Collembolas | Enchytraeids | Lumbricidae | Acari | Symphyla | Julida |
| --- | --- | --- | --- | --- | --- | --- | --- |
|  | Total annotator count | 558 | 1330 | 27 | 311 | 819 | 62 |
| ConvDirect | Total estimated count | 594 | 355 | 23 | 254 | 1120 | 308 |
|  | Rel. pred difference % (total) | 7 | 73 | 15 | 18 | 37 | 397 |
|  | Rel. pred difference % (7d) | 47 | 73 | 59 | 24 | 49 | 393 |
| ConvBinMul | Total estimated count | 504 | 595 | 25 | 258 | 1093 | 468 |
|  | Rel. pred difference % (total) | 10 | 55 | 7 | 17 | 34 | 654 |
|  | Rel. pred difference % (7d) | 41 | 57 | 52 | 26 | 46 | 650 |

### 3.3 Temporal dynamic of population activity according to the land-use context

Temporal population stability differed markedly between the treeline area (Position A) and the cultivated zone (Position C) across the majority of soil invertebrate taxa (Figure 4). Stability, quantified as the inverse of the temporal coefficient of variation (CV), was significantly lower in the cultivated zone for seven of the nine taxa examined. The strongest responses were observed for Lumbricidae (estimate = 1.24 *±* 0.06), Gastropoda (1.12 *±* 0.08), Acari (0.71 *±* 0.05), and Pauropoda (0.70 *±* 0.06), all showing substantially elevated CV in position C (log-link scale estimates *>* 0.5). Note that the Lumbricidae estimate was obtained from a random-intercept model, as the full AR1 model did not converge for this taxon; the directional conclusion is nonetheless supported by the congruence analysis below. Collembola (0.32*±*0.04), Insecta larvae (0.32*±*0.07), and Enchytraeidae (0.31 *±* 0.03) exhibited a similar but more moderate trend toward reduced stability in the cultivated zone. Symphyla was the sole taxon displaying the opposite pattern, with significantly higher stability in the cultivated zone (estimate = *−*1.10*±*0.06), indicating markedly lower CV in position C relative to the tree line area. Julida was the only taxon for which land use had no statistically significant effect on temporal stability, with a confidence interval broadly overlapping zero (estimate = 0.11 *±* 0.08; *p* = 0.18).

**Figure 4:**
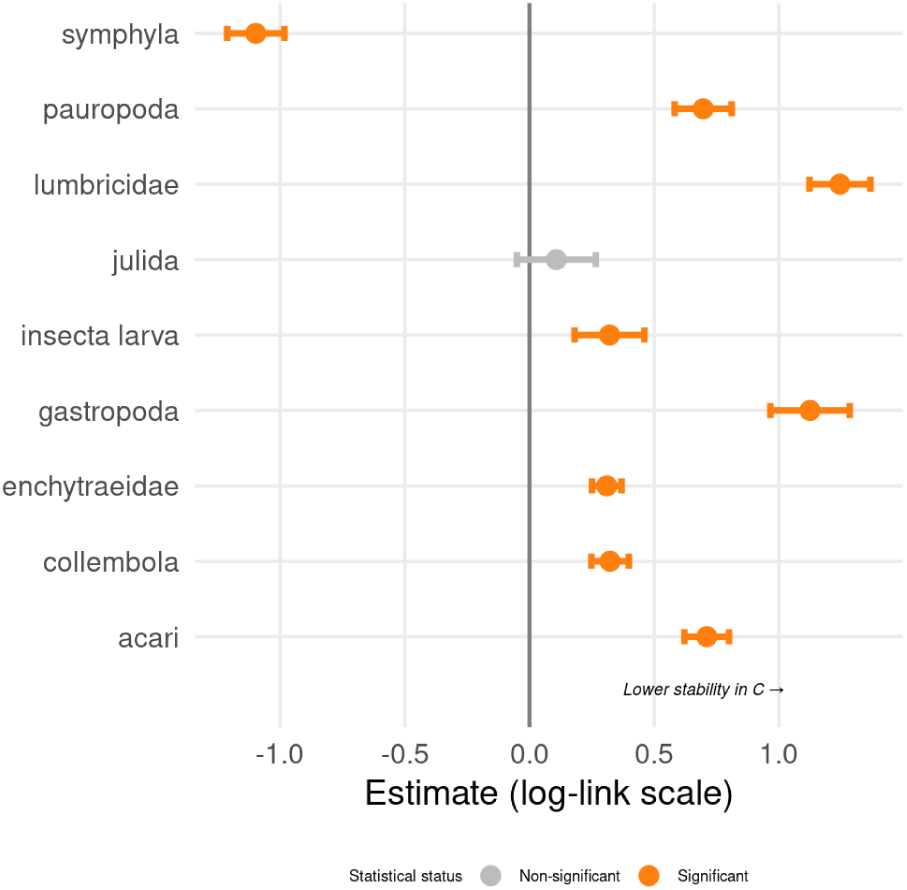
Points represent maximum-likelihood estimates of the Position C coefficient (cultivated zone versus arboreal linear feature) from Zero-Inflated Gamma generalised linear mixed models (log-link scale); horizontal bars span 95% Wald confidence intervals (estimate *±* 1.96 × SE). Positive values indicate higher temporal variability (lower stability) in the cultivated zone relative to the arboreal linear feature; negative values indicate the reverse. Taxa are ordered by decreasing effect size. Statistically significant effects (p < 0.05) are shown in orange; the non-significant taxon (Julida) is shown in grey. The Lumbricidae estimate was obtained from a random-intercept model (no AR1 structure) owing to non-convergence of the full model; all other estimates are from Zero-Inflated Gamma GLMMs with a first-order autoregressive correlation structure fitted over scanner orientation

Land-use effect sizes derived from the automated pipeline were largely congruent with those from expert-validated manual classifications (Figure 5), with all taxa clustering close to the 1:1 identity line. Gastropoda and Julida showed near-perfect agreement (*|*Δ*| ≤* 0.009 and 0.003, respectively), and Collembola also showed strong agreement (Δ = 0.036). Moderate discrepancies were observed for Enchytraeidae (Δ = 0.111), Acari (Δ = 0.143), Lumbricidae (Δ = 0.152), and Symphyla (Δ = 0.114), in all cases with the automated classifier overestimating the absolute effect size relative to human observers -with the exception of Lumbricidae and Pauropoda, for which the model underestimated the effect. Pauropoda exhibited the largest discrepancy overall (Δ = 0.246), while Insecta larvae also showed notable overestimation (Δ = 0.229). Therefore, despite classification errors, major ecological patterns (e.g. land-use effects on temporal variability) were consistently recovered.

**Figure 5:**
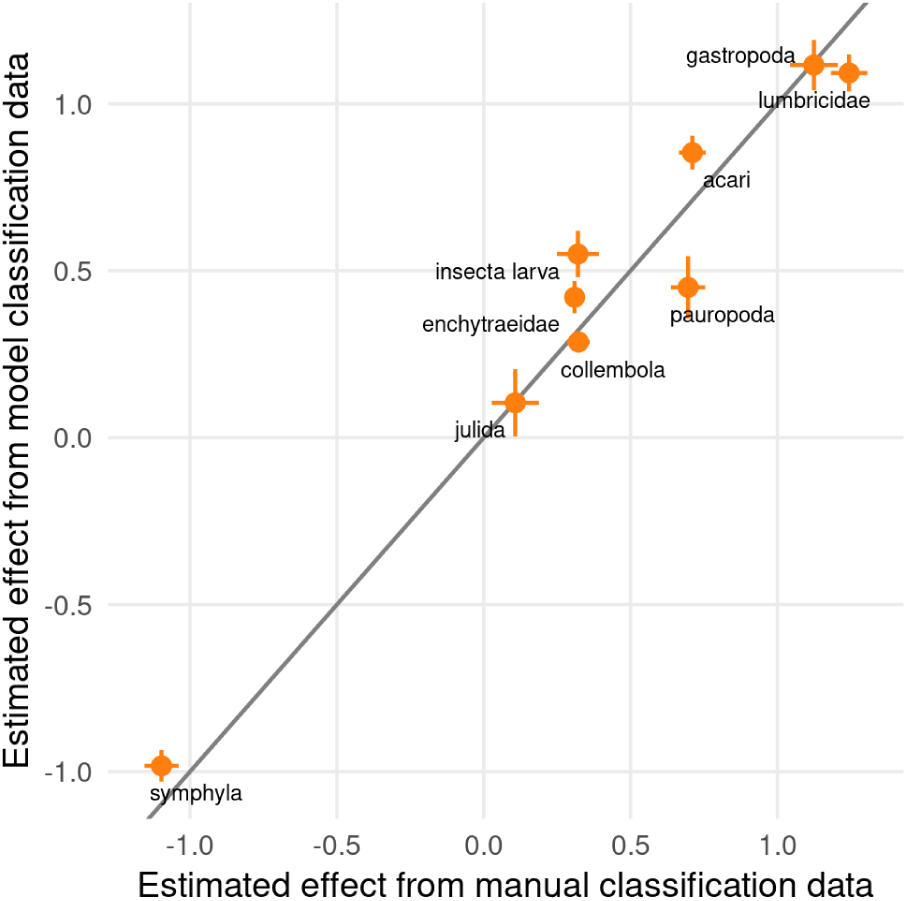
Technological congruence between manual expert classification and automated computer vision pipeline for the estimation of land-use effects on soil invertebrate population stability. Each point represents one taxon; the x-axis shows the Position C effect size (cultivated versus arboreal) estimated from expert-validated manual classifications, and the y-axis shows the corresponding estimate from the automated pipeline. The grey dashed line represents perfect 1:1 agreement. Error bars span *±* 1 SE on both axes. Taxa plotting close to the identity line show high cross-pipeline congruence. Effect sizes are on the log-link scale and are directly comparable across taxa

## 4 Discussion

### 4.1 From automated sensing to ecological inference

Recent advances in sensing technologies and machine learning have shifted the main bottleneck of biodiversity monitoring from data acquisition to data interpretation [Besson et al., 2022, Kitzes et al., 2026, Tuia et al., 2026]. High-frequency, continuous ecological datasets can now be generated at previously unimaginable scales, yet extracting biologically meaningful signals from them remains a fundamental challenge. This is especially true in soil, where the raw data is structurally complex, the signal-to-noise ratio is low, and annotated training resources are scarce. The present study demonstrates that this challenge can be substantially overcome through a hybrid pipeline combining heuristic detection with fine-tuned deep learning classification, enabling continuous soil fauna monitoring at spatio-temporal resolutions previously inaccessible to ecologists.

We argue that automated pipelines should not be viewed merely as technical tools, but as observation processes in their own rigth – each step of which introducing specific forms of uncertainty that shaped the resulting ecological metrics. In our case, detection based on image differencing achieved high recall but introduced false positives, while classification performance varied markedly across taxa (Tables 2 and 3), and the aggregation of detections into counts required calibration choices that influenced temporal variability estimates. Automated monitoring does not eliminate observation bias; rather, it redistributes it within the data-processing workflow.

Our work required sustained interdisciplinary collaboration from the outset – not as a sequential division of labour in which ecologists produced data and subsequently commissioned computational analysis, but as genuine co-design of the full analytical workflow [Carey et al., 2019]. The pipeline decisions that most determined final ecological performance – framing detection as a difference-based classification problem, constructing scanner-aware cross-validation folds to prevent data leakage, and selecting population counts rather than classification accuracy as the primary validation metric – each demanded simultaneous ecological and computational expertise. Rigorous quality control was similarly impossible without the combined contributions of model developers, who could identify systematic failure modes, and domain practitioners, who possessed the local and taxonomic knowledge needed to interpret them [Tuia et al., 2022]. The possibilities for advancing ecological understanding through emerging data streams are immense, but realising them requires multidisciplinary frameworks that treat data collection and data analysis as inseparable components of a single scientific process [McCrea et al., 2023].

### 4.2 Pipeline choices and future directions

We now discuss several technical aspects related to the pipeline.

Recall of the heuristic detection algorithm. The heuristic detection step prioritizes sensitivity, ensuring that most organisms present in the images are captured. A key question for the pipeline’s validity is how many individuals are lost at this stage – a quantity that is intrinsically difficult to evaluate since the ground truth requires annotating raw scans in their entirety. The cross-site count comparison (Table 3) does not reveal systematic underestimation patterns for most taxa, suggesting that the high-recall configuration of the detection stage performs as intended – which also comes with a significant number of false positives, including noise and non-biological movements. Conversely, false negatives were more likely to occur for low-mobility or poorly contrasted organisms. The exception is Enchytraeidae, whose low count accuracy is more plausibly attributed to classification error than to detection failure – as discussed later in this section. Dedicated recall validation on a representative image subset remains an important objective for future work, which may be addressed by comparing results with a different automated method, to better understand the biases of our observation process. Yet, this low-precision high-recall stage is only beneficial if false detections can be identified efficiently, which we now discuss.

Binary classification model. The dedicated binary model Binary achieved high micro-averaged mAP, confirming its effectiveness at discarding the bulk of background boxes. However, its substantially lower binary macro-averaged mAP of 67% – relative to the scores produced by the multiclass classifier ConvDirectof 86% – reveals that its discriminative capacity concentrated on frequent invertebrate classes rather than generalising across all taxa. ConvDirect, by contrast, produced considerably higher binary macro-averaged mAP, suggesting that taxon-specific training objectives produce features that are more broadly generalisable to the invertebrate-versus-background distinction than a binary training objective alone. This may also be due to the increased model capacity – there is one output logit for each class, since binary macro-averaged mAP performances drop if all invertebrates scores are aggregated into a single class. This disparity in classification accuracy across taxa questions the relevance of binary prefiltering for our problem.

Direct multiclass classification versus two-stage model. The two-stage classification architecture ConvBinMul – in which a binary pre-filter is followed by a dedicated multiclass classifier – achieved marginally higher average performance than the direct multiclass model ConvDirect, but introduced three sources of additional complexity: it requires training and deploying two models rather than one; the training dataset must be pre-filtered to discard background boxes at an appropriate threshold; and the binary rejection threshold must be tuned prior to the multiclass step. This tuning is sensitive to the data distribution and can substantially affect final count estimates, as evidenced by the contrasting cross-site behaviours of the two architectures (Table 3). Because this sensitivity demands careful calibration, as it risks degrading generalisation performance under distributional shift – precisely the scenario most relevant to deployment, ConvDirect was adopted as the default pipeline, notwithstanding the aggregate mAP advantage of ConvBinMulwhen adequate tuning is possible. Interestingly, despite slight differences in average performance, hard and easy classes seem to be the same for both approaches, indicating for instance that binary prefiltering does not affect binary mAP as we compute it in a substantial way.

Multiclass classification performances. Classification performance varied markedly across taxa (Tables 2 and 3), with some groups consistently achieving high accuracy while others remained difficult to distinguish. Macro-averaged mAP and macro-averaged binary mAP are computed using the same model scores, but differ in the composition of the evaluation set: in the binary setting, samples from classes other than the focal taxon and background are excluded, whereas all classes are retained in the multiclass one-versus-all setting. Comparing results across these two settings reveals two distinct classification difficulty profiles among the invertebrate taxa. The first comprises taxa that are difficult to detect against background but straigthforward to classify once detected - including Acari, Collembola and Gastropoda, for which multiclass mAP and binary mAP are closely aligned. The second comprises taxa that are reliably detected but difficult to assign precisely: Pauropoda (binary mAP 88%, multiclass mAP 73%), Julida (binary mAP 96%, multiclass mAP 80%) or Lumbricidae exhibit the largest gaps between these two metrics. Importantly, these differences did not result in uniform noise, but in structured biases: taxa with lower performance contributed disproportionately to variability in estimated counts, whereas well-classified taxa provided stable signals (Figure 3). Similarly, taxa that are not reliably detected risk under- (or over-) detection, and are thus much more sensitive to the number of false positives or changes in the background (which may complicate detection even further). Note that while we do believe we provide a reasonable baseline, performance improvements can surely be obtained by testing more architectures, hyperparameters, and training strategies. Yet, our goal at this stage is not to gain a few percentages of mAP, but to validate the relevance of the workflow, which includes high-level design choices – e.g., high-recall low-precision heuristic detection followed by efficient classification – or defining the right metrics – which depend on the downstream ecological tasks. We leave further optimization of individual pipeline blocks for future work.

Training set size and classification accuracy. We identify an expected correlation between the number of annotated examples per taxon and relative model count errors. Yet, Figure 3 still features variability, indicating that taxon identity – and by implication the morphological and behavioural characteristics that determine detection and classification difficulty – plays a key role in pipeline accuracy. Enchytraeidae illustrate this clearly: despite comprising the second-largest annotated class, they exhibited the highest count errors of all taxa, likely due to their white elongated transparent morphology similar to roots (see Appendix A). Symphyla, by contrast, achieved low count errors despite being comparatively rare. While expanding training data will always be beneficial, targeted methodological approaches – such as integrating short temporal sequences to exploit movement kinematics, or morphology-specific augmentations – are likely to yield greater gains than additional annotation alone for persistently difficult taxa. Beyond improving the classification scores they rely on, the population counts accuracy may be improved by changing how we convert scores into meaningful counts.

From classifier scores to population counts. The aggregation of detections into ecological metrics (e.g., population counts) introduced an additional layer of transformation. We derived population counts by assigning each candidate box to the highest-scoring taxon class. Preliminary experiments with alternative count estimation strategies that leverage the full classifier scores distribution did not improve performance, suggesting that the current hard-assignment approach is a very strong baseline at this dataset scale. A promising complementary direction would be to compensate for systematic over- or under-counting of specific taxa using site-level calibration statistics, as is for instance standard in remote sensing [Olofsson et al., 2014]. An complementary approach would be to borrow from conformal prediction, and define taxon-specific thresholds [Fontana et al., 2023, Leblanc et al., 2025]. However, the opposing directions of Enchytraeidae estimation error at the DIAMs site (overestimated, Table 2) and the external truffle-oak site (underestimated, Table 3) demonstrate that any such correction would be site-specific and may actively degrade cross-site transferability. Improving generalisation across soil types, geographic regions, and invertebrate community compositions therefore remains the principal open challenge for future pipeline development.

Generalisation in a technical sense, however, is only one dimension of pipeline validity. Our results demonstrate that automated pipelines define a composite observation model, where multiple sources of uncertainty – detection errors, taxon-specific classification performance, and aggregation choices – interact to shape ecological outputs. The more demanding test is whether the ecological conclusions the pipeline generates can be trusted – that is, whether the biological narrative it produces is consistent with that established by expert human observers.

### 4.3 Ecological validity and robustness of inference

A central question for automated monitoring is whether ecological conclusions remain valid when derived from model-generated rather than expert-based observations. Our results provide strong support for this, showing that major ecological patterns are preserved despite imperfect classification.

The land-use contrast between arboreal linear features and cultivated zones produced a consistent and biologically interpretable pattern across eight of nine taxa, with cultivated zones exhibiting substantially higher temporal variability in population activity. This finding aligns with the established role of tree strips in agroforestry systems as structural refugia [Marsden et al., 2020]: the structural continuity of the arboreal feature provides stable microhabitat conditions – moisture buffering, reduced mechanical disturbance, persistent litter inputs – that buffer invertebrate communities against the pulsed dynamics imposed by tillage and crop cycles in the adjacent cultivated soil. The single exception, Symphyla, showed markedly higher stability under cultivation – a counterintuitive results that may reflect this taxon’s specific sensitivity to intensified biotic interactions, as competitive pressure or predation, associated with the more diverse and active community of the arboreal zone. The mechanisms underlying this response warrant dedicated investigation, but its detection by both the manual and automated pipelines independently strengthens confidence in its biological reality.

The agreement between automated and expert-derived estimates of land-use effects was high across all nine taxa (Figure 5), and critically, the relationship between classification performance and inference reliability was structured and predictable: taxa with high mAP contributed stable, congruent signals across methods, while taxa with lower performance showed greater discrepancies in estimated effect sizes. This proportionality is reassuring precisely because it means that pipeline limitations are traceable rather than arbitrary —-users can anticipate where automated inference is most and least reliable based on independently measurable classification metrics. Similar conclusions have been reached in other domains of automated biodiversity monitoring, where ecological signals can remain robust to imperfect detection and classification [Besson et al., 2022, Weinstein, 2018]. Our results reinforce this principle and extend it to a novel environment and data type, while providing an empirical framework for defining the taxon-specific boundaries of reliable inference.

More generally, these findings suggest that a key criterion for evaluating automated pipelines is not absolute classification accuracy, but the preservation of ecological inference. While both are linked to a certain extent, moderate classification performance was sufficient to recover consistent ecological patterns in our case. This reframing has practical implications: it shifts the validation priority from optimising benchmark metrics towards explicitly testing whether the ecological conclusions of interest are robust to the specific error structure of the pipeline being deployed.

Finally, the 33-day time series analysed here represents only a fraction of the ecological signal that the automated pipeline makes accessible. Manual annotation at this temporal scale was feasible but required sustained expert effort that cannot be scaled to multiple sites or seasons. Extending the same analytical workflow to a full annual cycle – encompassing seasonal transitions, agricultural management events, and inter-annual variability – would be entirely impractical without automated classification. This contrast between what is possible with and without the pipeline constitutes the clearest argument for its ecological value: it does not merely replicate what human annotation achieves; it enables study designs that human annotation cannot.

### 4.4 Toward generalizable and reproducible automated monitoring workflows

Beyond the specific case study presented here, this work contributes to the development of generalizable workflows for automated biodiversity monitoring. The release of our annotated dataset – comprising over 8,000 labeled invertebrate detections across ten taxa – represents the largest publicly available resource of its kind for in situ soil fauna monitoring to our knowledge. In a nascent field, the proliferation of isolated, incompatible datasets risks fragmenting progress. By deliberately providing these data and our processing pipeline as open infrastructure – a practice not yet standard in the field – we aim to facilitate the benchmarking and community-wide standardization – covering annotation protocols and taxonomic resolution – necessary for the large-scale, multi-site analyses required to understand biodiversity at continental scales [Tuia et al., 2026, Kitzes et al., 2026].

However, our results also highlight key challenges for generalization. Model performance and calibration varied across taxa, scanners, and evaluation settings (Tables 2 and 3), indicating that models trained under specific conditions may not directly transfer to new ecological contexts. Addressing these challenges will require the development of standardized datasets and evaluation protocols that explicitly account for ecological variability [Besson et al., 2022].

A final critical frontier is the formal integration of pipeline uncertainty into statistical inference. While our study demonstrates that automated pipelines can recover major ecological signals (e.g., land-use effects), we do not yet explicitly model the propagation of false positives and misclassifications into final population estimates. Future work should prioritize the development of observation models that treat the automated pipeline as a known source of uncertainty [Kitzes et al., 2026]. Ultimately, Fully leveraging automated sensing technologies will require not only technical advances, but also conceptual frameworks that integrate observation processes into ecological inference. By bridging the gap between computer vision and ecological modeling, we can ensure that the increasing availability of high-resolution data translates into reliable ecological understanding [Weinstein, 2018, Besson et al., 2022].

## 5 Conclusion

We have presented and validated an end-to-end automated pipeline for converting raw image sequences into taxon-resolved population counts. The pipeline – combining a data-free heuristic detection stage with a fine-tuned deep learning classifier – substantially reduces the annotation burden that has previously constrained monitoring based on image time-series to short time windows and limited taxonomic scope, while preserving the ecological conclusions established by expert human classification across all nine taxa examined. The accompanying annotated dataset, the largest of its kind for in situ soil invertebrate imaging, provides a community resource for benchmarking, transfer learning, and methodological extension.

Our pipeline was designed for counting invertebrates from soil optical imaging data, and we demonstrated its ecological validity through a comparative stability analysis across contrasting land-use contexts. Thus, our work represents a concrete step towards routine, high-resolution monitoring of soil fauna communities at temporal and spatial scales commensurate with the ecological questions – seasonal dynamics, land-use impacts, biodiversity trends – that the discipline most urgently needs to address. Realising this potential more broadly will require continued interdisciplinary collaboration, coordinated data sharing, and systematic validation workflows that hold automated outputs to the same standard of ecological rigour as manually derived data.

## A Additional information about the dataset

The dataset we release can be accessed by following this link. Note that the boxes were extracted using a previous R prototype of the heuristic detection algorithm, which does not exactly correspond to the Python version with release. We release the R code as supplementary material for full reproducibility, but also provide the Python code so that the full workflow does not require two separate programming languages.

### A.1 Metadata

#### A.1.1 Raw scans

Images were collected at a 6-hour temporal frequency from March 6 to May 19, 2024 in A4 format (21×29.7cm) at 1200 dpi, leading to 590 RGB images of size 10200×14032, stored in jp2 format.

#### A.1.2 Image extracts

Boxes are split among 13 classes, with varying number of images per class: acari (291), collembola (2837), formicidae (58), insecta larva (66), lumbricidae (445), pauropoda (67), background (47814), enchytraeidae (930), gastropoda (634), julida (103), multi_taxa (324), symphyla (363), and unknown (1881).

Statistics about the size of boxes that contain invertebrates and number of detections per scan are shown in Figure 8. They illustrating strong variability in terms of area, with a relatively consistent number of detections per image. Images are stored in jpeg format.

**Figure 6:**
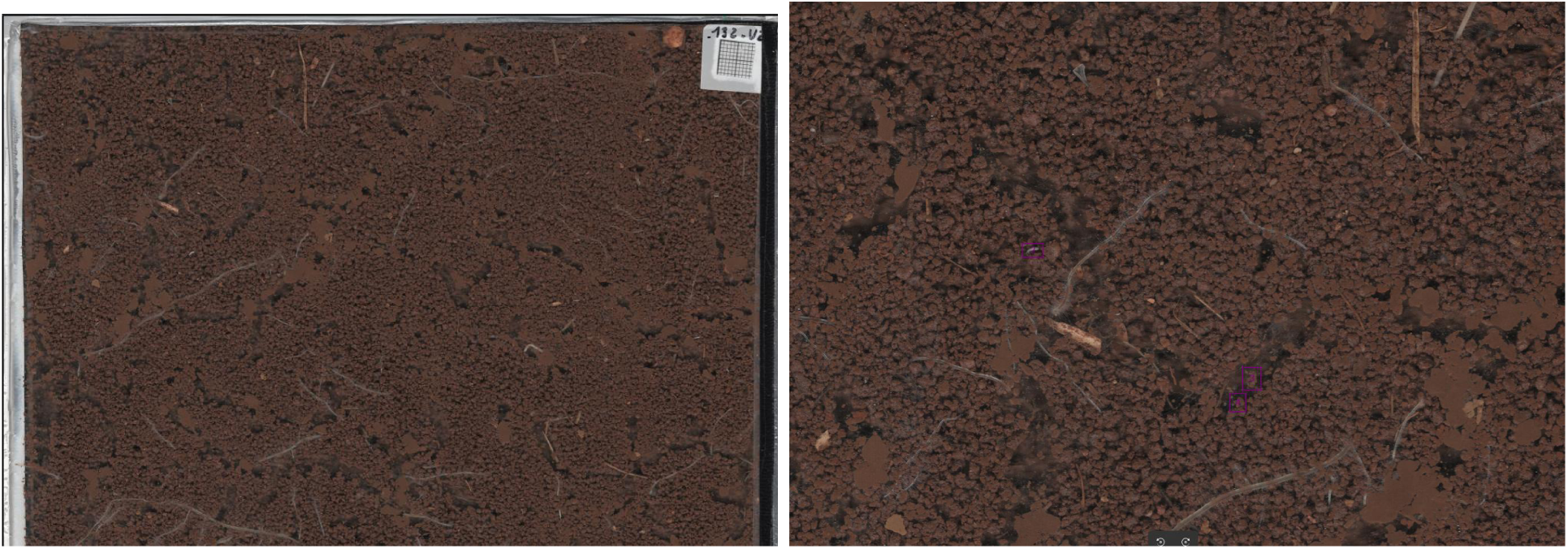
Sample images from the original dataset. Left: top half of a raw scan image. Right: Zoom on a smaller image region, with candidate detection boxes (in purple).

### A.2 More example images

We display here a few images from the different stages of the pipeline. Figure 6 shows the raw input data, i.e., the large scans, as well as results of our heuristic detection pipeline on part of the base image. Then, 7 gives five different examples for each class of invertebrates.

### A.3 Consolidating ground-truth labels

An audit of the 300 boxes assigned the highest invertebrate scores despite being labelled as background (top-scoring false positives) identified approximately 100 incorrectly labelled invertebrate individuals — an overall error rate below 2%, yet revealing a strongly directional annotation bias. No reciprocal errors were found among invertebrate boxes assigned high background scores, confirming that label noise is asymmetric and driven primarily by fatigue: because background boxes dominate the dataset, annotators operating under sustained workload tend to default to this class, selectively missing invertebrates rather than generating false positives. A score-guided annotation strategy — prioritising high-scoring boxes for review — could reduce this bias and improve label quality, particularly for taxa such as Pauropoda that are reliably detected but poorly classified. However, such a strategy would simultaneously reinforce biases for poorly-detected taxa such as Enchytraeidae, whose hard examples are precisely those the model already tends to miss. Any iterative workflow in which model outputs guide annotation must therefore be implemented with care to avoid the progressive self-reinforcement of model errors [Desprez et al., 2023].

**Figure 7:**
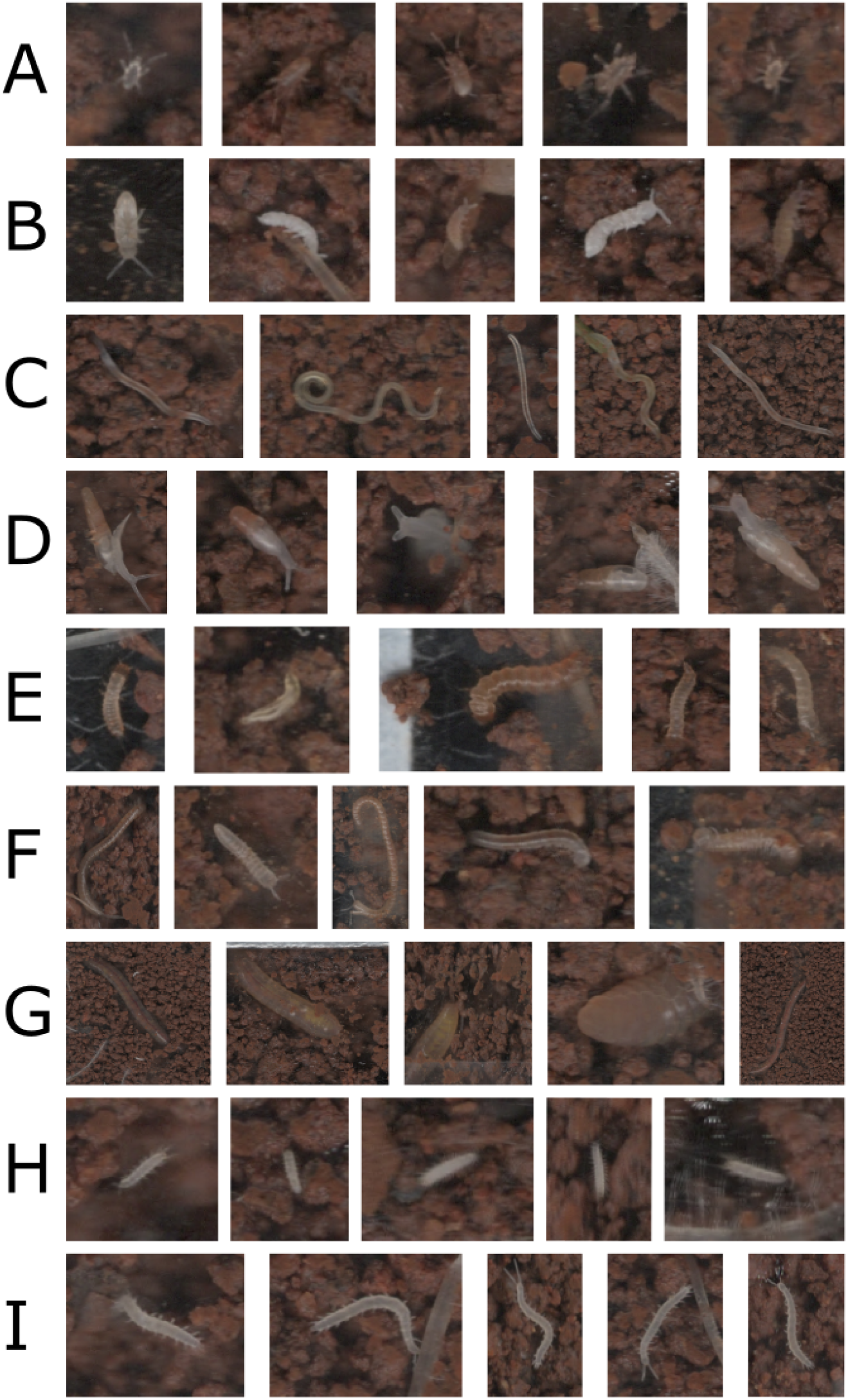
Five images per taxa, illustrating the variability of invertebrates that need to be identified. All images have been resized. A : acari; B : collembola; C : enchytraeidae; D: gastropoda; E: insecta larva; F: julida; G: lumbricidae; H: pauropoda; I: symphyla

**Figure 8:**
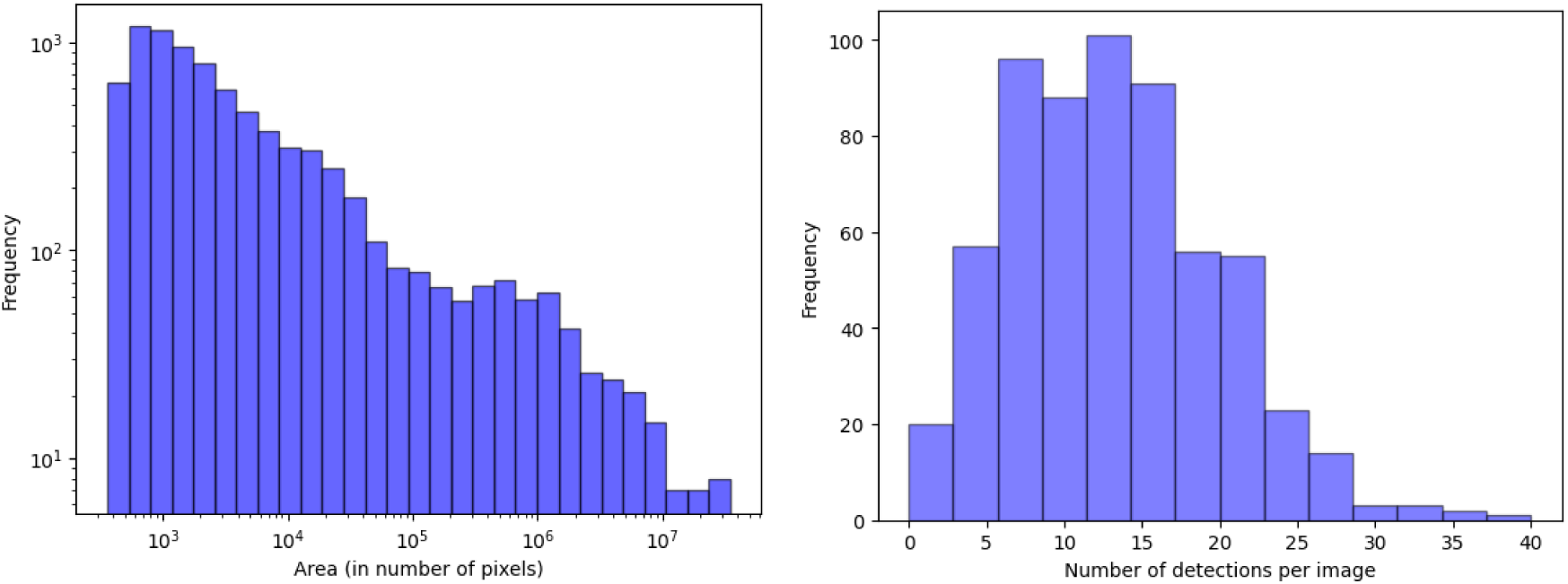
Left: Histogram of invertebrates boxes area. Right: Histogram of invertebrates detections per scanner image. The boxes are obtained using the heuristic detection pipeline described in Section 2.2, which leverages differences of successive images. We see that many boxes contain at most a few thousand pixels, which are extracted from base scans with 1.4 *×* 10^8^ pixels, i.e., 10^5^ times bigger.

## Footnotes

1 The dataset can be found following this link.

2 Our code can be found here: https://gitlab.com/f.postic/deepsoilfauna

## Notes

https://gitlab.com/f.postic/deepsoilfauna

https://zenodo.org/uploads/20662250

### Competing Interest Statement

The authors have declared no competing interest.

### Funding

- Agence de la transition écologique: CASCA

